# Unconstrained multivariate EEG decoding can help detect lexical-semantic processing in individual children

**DOI:** 10.1101/797175

**Authors:** Selene Petit, Nicholas A. Badcock, Tijl Grootswagers, Alexandra Woolgar

**Author notes:** Corresponding author at: Department of Cognitive Science, Macquarie University, Australia.

## Abstract

In conditions such as minimally-verbal autism, standard assessments of language comprehension are often unreliable. Given the known heterogeneity within the autistic population, it is crucial to design tests of semantic comprehension that are sensitive in individuals. Recent efforts to develop neural signals of language comprehension have focused on the N400, a robust marker of lexical-semantic violation at the group level. However, homogeneity of response in individual neurotypical children has not been established. Here, we presented 20 neurotypical children with congruent and incongruent visual animations and spoken sentences while measuring their neural response using EEG. Despite robust group-level responses, we found high inter-individual variability in response to lexico-semantic anomalies. To overcome this, we analysed our data using temporally and spatially unconstrained MVPA, supplemented by descriptive analyses to examine the timecourse, topography, and strength of the effect. Our results show that neurotypical children exhibit heterogenous responses to lexical-semantic violation, implying that any application to heterogenous disorders such as ASD will require individual-subject analyses that are robust to variation in topology and timecourse of neural responses.

## 1 Introduction

In a variety of conditions, the absence of spoken language does not necessarily reflect an absence of comprehension. In addition, poor language production is not always associated with low verbal or non-verbal IQ ^1–3^. For populations that are unable to reliably communicate, such as people in a vegetative state ^4^, or with cerebral palsy ^5^ or non-verbal autism ^6^, an accurate assessment of cognitive and language aptitudes is essential, but challenging. Traditional testing materials often fail to capture the full cognitive and language abilities of these individuals. This is due to the various constraints imposed by the standardised testing environment and materials, and the social constraints associated with the examiner-examinee interactions ^7,8^. To assess cognitive and language abilities of these populations, we need to develop objective, fast, and reliable measures of cognition and language comprehension. Neuroimaging (e.g., electroencephalography – EEG, functional magnetic resonance imaging – fMRI) allows the indirect observation of neural processing, bypassing the need for behavioural responses. For example, FMRI has previously been used to assess cognition in non-communicative populations such as patients with disorders of consciousness ^9,10^. EEG is a less expensive and more readily alternative which may also be suitable for passively measuring cognitive responses in the absence of reliable behaviour. Recent work has used EEG to assess covert cognitive abilities in non-communicative populations (see a recent review by Harrison and Connolly ^11^). In particular, language processing has been studied using EEG including in patients with disorder of consciousness ^12,13^, schizophrenia ^14,15^, and autistic individuals ^16–18^[Note : we use ‘identify-first’ language (‘autistic person’) rather than person-first language (‘person with autism’), because it is the preferred term of autistic activists (e.g. Sinclair, 2013) and many autistic people and their families ^20^ and is less associated with stigma ^21^]. However, in these cases, EEG research has focused on the population-level, with minimal data reported on an individual-participant basis. Yet, in order to design a clinical test of language comprehension, and particularly given the known heterogeneity in developmental disorders such as ASD, it is critical to use paradigms and methods that reliably elicit meaningful brain signals in individuals. In this study, we therefore developed and assessed a new EEG paradigm to measure language comprehension in individual children. We report the heterogeneity of neural responses to semantic anomalies of speech using different data analysis techniques.

The N400 ERP is a well-studied component that indexes the semantic integration of words into their context. The N400 effect is a more negative-going electrical potential when participants read or listen to words presented in an incongruent semantic context, relative to a congruent semantic context ^22^. As such, it seems to be a well-suited candidate for the assessment of lexico-semantic processing. The N400 has been widely reported in groups of typical adults, children, and special populations, including recently in autistic children ^23–25^. However, despite the recent interest in using N400 as an assessment tool, individual subject level analyses are rarely reported. Additionally, studies that do report individual-subject data are often limited by methodological or conceptual factors, such as a lack of proper statistical analyses, or lack of a control group. Recently, two studies have looked at the individual sensitivity of auditory N400 paradigms, in healthy adults ^26^, and in healthy children ^27^. These studies each used two paradigms contrasting auditory words presented in congruent or incongruent semantic contexts. Using a task in which participants were asked to make covert violation decisions, these studies reported a statistically significant N400 effect in a maximum of 58% of neurotypical adults ^26^ and 56% of neurotypical children ^27^. This moderate sensitivity reflects the challenges associated with assessing ERPs at the individual level. However, using multivariate pattern analyses (MVPA) on these EEG data, we recently increased the individual-subject detection rate to 88% ^27^. In the current study, we aimed to build on existing N400 paradigms by improving the stimulus delivery paradigm and by using a more robust and sensitive analysis framework. Our goal was to derive a highly sensitive measure of language comprehension in individual children.

We set out to extend the paradigm developed in ^27^, and to measure neural differential responses to auditory words presented in congruent and incongruent sentences frames (i.e., “The squirrel stored nuts in the *tree/door*”). In order to increase children’s engagement to the task, and build up strong semantic contexts, the spoken sentences were accompanied by short animated cartoons (*e.g.*, an animation of a squirrel storing nuts in a tree). With clinical applications in mind, our paradigm presented a semi-covert task, in which participants were asked to silently judge the semantic congruency of each sentence in their head, with occasional button press requests to check for compliance. Additionally, we explored several analyses of the EEG data to illustrate the inter-individual variability of neural responses.

With traditional within-individual ERP analyses, it is necessary to choose *a priori* time windows, and electrodes, of interest, in order to reduce the number of comparisons and thus increase the statistical power. However, we have previously demonstrated that for individual subject analysis, *a priori* assumptions about spatio-temporal location should be avoided as there is substantial inter-individual variability in the location and timing of N400-like effects ^27^. For clinical populations, for whom inter-individual variability may be even higher, it is essential to allow some inter-individual variability in the location and timing of the effect of interest. To allow for this without increasing multiple comparisons, we previously used multivariate pattern analyses (MVPA). This approach uses the signal recorded across all the EEG channels to detect patterns of brain activation that reliably distinguish between two conditions, in this case, between identical words presented in different lexico-semantic contexts. Using MVPA at each time point, we retain information about when an effect occurs, but allow it to arise from any spatial location. In the current study, we again used MVPA, but additionally dropped the requirement for time-resolved results, allowing the classifier to detect an effect with any spatial configuration and any temporal profile in an effort to increase our detection rate. This allowed us to detect differences in the brain’s pattern of activity irrespective of the location and the timing of the difference. Having detected a statistical difference using this approach, time-resolved MVPA and univariate approaches can then be used to qualitatively describe the temporal and topographic distribution of the effect. In this study, we sought to design an engaging, covert paradigm to elicit individuals’ neural responses to semantic violations within spoken sentences. We contrasted congruent and incongruent sentences, and measured the brain response of children using EEG, and we analysed the data using MVPA and univariate N400 analyses. We found the strongest detection rate using an unconstrained MVPA approach, reaching a medium detection rate (13/20 participants). Univariate analyses were less sensitive, possibly due to the need to restrict the analyses to a predefined brain region for appropriate statistical power. We additionally provide evidence for this claim by illustrating the high inter-individual variability in the location and timing of the discriminative neural signals.

## 2 Methods

All presentation scripts, stimuli, and raw data available at https://osf.io/bv2dy/.

### 2.1 Stimulus development and validation

#### 2.1.1 Congruent stimuli

The stimuli consisted of 94 congruent and 94 incongruent sentences. The congruent sentences were adapted from the norms of ^28^, who reported the cloze-probability of 498 sentences for adults in the USA. We started by selecting the 450 sentences with a cloze probability higher than 50%. (*i.e.*, more than 50% of the participants completed a given incomplete sentence with the same target word), and for which the target word was a noun. From this set, we then selected only the 242 sentences in which the target word and of all the keywords were of high frequency (Zipf Log10 frequency > 3.5), according to the children section of the SUBTLEX-UK word database ^29^. This removed sentences that were not suitable for children.

To facilitate the splicing of the target word from the audio recording, we retained only the 105 sentences in which the boundary between the incomplete sentences and the target was a plosive sound (/t/, /d/, /k/, /p/, /b/, or /g/), and the 35 sentences in which it was possible to add an adjective ending in a plosive sound before the target without disrupting the meaning of the sentence (*e.g.*, To cut the chicken Sue needed a *sharp* knife – see Table 1). The remaining 140 sentences were recorded by a female, native Australian-English speaker. In order to make sure that the congruent sentences were highly congruent for children, eighteen native English speaker children, aged 8- to 12-years old (M = 9:11, range = [8:4 to 11:11]) participated in a validation experiment. We presented participants with each of the 140 incomplete sentences, and asked them to say the word they thought would best complete each sentence. We then selected only the sentences with a target cloze probability of over 60%, leaving a full stimulus set of 94 sentences (see Supplementary Table S1 for a complete list of the stimuli). The mean length of the sentences was 3.78 s (SD = .68 s, range = [2.40; 6.21]).

**Table 1.** Example of congruent and incongruent sentences. The conditions are matched by pairing the targets between the two conditions. In some cases (last two rows), we added an adjective (italics) ending with a plosive sound before the target word to facilitate audio splicing (see Supplementary Table S1 for a complete list of the stimuli).

| **Incongruent sentence** | **Congruent sentence** | **Target** |
| --- | --- | --- |
| In autumn, leaves fall off the | After dinner, they washed the | dishes |
| After dinner, they washed the | In autumn, leaves fall off the | trees |
| They raised pigs on their *private* | To cut the chicken Sue needed a *sharp* | knife |
| To cut the chicken Sue needed a *sharp* | They raised pigs on their *private* | farm |

#### 2.1.2 Incongruent stimuli

In order to generate an incongruent condition in which the target words and sentence frames were perfectly matched with the congruent condition, we swapped the target words of pairs of congruent sentences (see Table 1). Each target and each incomplete sentence was thus presented once in the congruent condition, and once in the incongruent condition. When recombining targets with incongruent sentences frames, we ensured that the target did not violate syntactic structure of the sentence, including matching for plurality. Furthermore, we ensured that the incongruent target did not start with the same sound nor rhyme with the congruent target.

### 2.2 Animations

For each congruent sentence, we designed a short, colourful, animated cartoon using that matched the meaning of the sentence, that were drawn and animated by Gabriella Keys using Adobe Photoshop CC 2017. The animation corresponded to the congruent version of the sentence, and each animation was presented twice, once in each condition (*e.g.*, the animation of a leaves falling off autumnal trees was presented with the sentence “in autumn, leaves fall off the *trees*”, and with “in autumn, leaves fall of the *dishes*”).

### 2.3 EEG experiment

#### 2.3.1 Participants

Twenty children aged 9 to 12 years (M = 10:6, SD = 0:11, 12 males and 8 females) were recruited through the Neuronauts database of the Australian Research Council Centre of Excellence in Cognition and its Disorders. All participants were native English speakers, and received $25 for their participation. This study was approved by the Macquarie University Human Research Ethics Committee (Reference number: 5201200658), and all methods were performed in accordance with the relevant guidelines and regulations. Informed consent was obtained from a parent and/or legal guardian for each participant.

#### 2.3.2 Experimental procedure

Participants were seated in front of a computer screen in a lightly dimmed room. The auditory sentences were presented via speakers on both sides of the screen, and the corresponding animations played on the screen at a visual angle of approximately 4° of height and width.

All 188 sentences were presented once within a single recording session, in a pseudo-random order that was optimised to minimise bias in the sequence of congruent and incongruent trials. For this, we generated 1000 candidate sequences, all constrained to have no more than 4 trials of the same condition in a row, and such that longer sequences of the same condition were no more frequent than shorter ones. We then selected the sequence that minimised first, second and third order bias in whether a given trial would be congruent given the preceding trials. Additionally, during the experimental session we constrained the order of the sentences so that each of the sentence frames was used once in the first half and once in the second half of the experiment. For example, the sentence frame “It was windy enough to fly a” was paired with the target word “kite” (congruent) and “tongue” (incongruent). Therefore, if the sentence “it was windy enough to fly a kite” was presented in the first half of the experiment, then the sentence “it was windy enough to fly a tongue” could only appear in the second half. We chose to present each complete sentence only once to minimise repetition effects and to limit the length of the recording session. For our main analysis (below) we calculated that 94 trials per condition would give us 96% power to detect a medium effect (Cohen’s d = 0.5) at the individual-subject level.

Each trial started with a central fixation cross, displayed for two seconds, followed by the presentation of a sentence. The animation started first, and included a 500 ms gradual fade-in and a 500 ms fade-out to minimise abrupt onsets and reduce eye-movements. After 500 ms, the auditory sentence started. To ensure that each animation was at least three seconds long irrespective of the auditory sentence length, we added a silent pause before each auditory sentence that was shorter than three seconds. The animation disappeared one second before the target word was presented, to ensure that EEG responses to visual information did not contaminate the response to the target word (see Figure 1).

**Figure 1.**
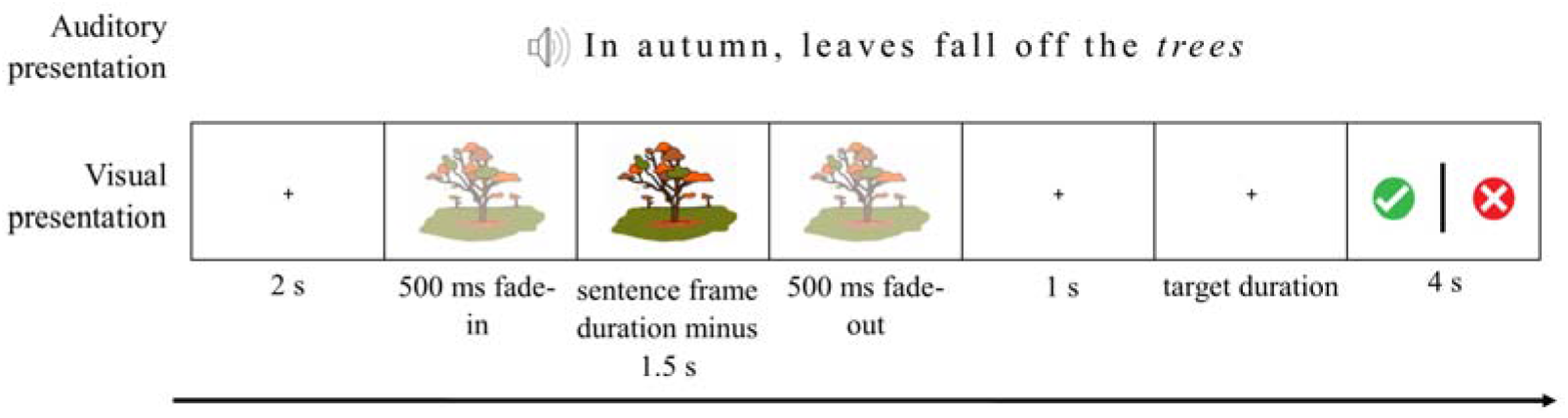
Trial sequence on a question trial. Note that the same animation would be presented for the incongruent version of this sentence (“in autumn, leaves fall of the *dishes*”). On non-question trials the final display was not shown and the 2s ITI started immediately. Images were drawn and digitised by Gabriella Keys.

We presented the participants with six practice trials at the start of the experiment to ensure they understood the instructions. Participants were instructed to listen carefully to each sentence and covertly decide whether the sentence was correct or incorrect. Additionally, on some trials, participants were asked to quickly indicate their judgement by pressing a button with their right hand. On these trials a response screen was presented with a green “tick” (“correct meaning”) and a red cross (“incorrect meaning”) on either side of the display. A central vertical bar decreasing in size indicated the time left to answer (4 seconds in total). Question trials occurred approximately every 7 trials, with a jitter of +− 3 trials so they were unpredictable. The red cross and the green tick appeared pseudo randomly on either the left or right hand side of the screen, so participants could not anticipate which button they would have to press. If they answered correctly, the participant was told that they had caught an “evil alien”. If they answered incorrectly, they were shown a semi-masked picture of the alien that they “did not catch”. After each question trial they were told how many evil aliens they had left to catch.

#### 2.3.3 EEG recording and pre-processing procedures

We acquired EEG data from a 64-channel ActiveTwo BioSemi (BioSemi, Amsterdam, Netherlands). The electrodes were organized according to the 10-20 system, with two electrodes placed on the left and right mastoids for offline referencing. Electro-oculogram generated from eye movements and eyeblinks was recorded using two facial electrodes, located at the outer canthus of and under the right eye, respectively. The data were digitized at 512 Hz with an anti-aliasing filter with 3 dB point at 104 Hz (fifth order sinc filter) with an online reference to the common mode sense (CMS), and all impedances were kept below 30kΩ.

We processed the data off-line using the EEGLAB toolbox in MATLAB ^30^. We first re-referenced the data to the average of the left and right mastoids, then used a bandpass filter between 0.1 and 40 Hz as recommended in ^31,32^. We then segmented the data into 1100ms epochs time-locked to the onset of the auditory target word (100ms pre-stimulus and 1000ms post-stimulus). To correct for eye-blinks and eye-movements, we ran an Independent Component Analysis (ICA). We removed the components with scalp distributions, time-courses and spectral contents indicative of eye blinks and eye movements. Finally, we applied a baseline correction to the epochs, and either saved all the epochs for subsequent MVPA analyses, or removed epochs with extreme values (+−200mV) for subsequent univariate analyses. For the purpose of illustrating the topography of the ERP across the scalp, we removed epochs that had extreme values in any channel. At this point, any channel that contributed to the rejection of more than 10% of the trials at this point was interpolated using spherical interpolation, and the preprocessing was done again using the interpolated channel(s) from the filtering stage. An average of 1.9 channels were interpolated per participant (range = 0-8). None of the channels in the region of interest (see below) had to be interpolated. For the purpose of analysing the N400 within our region of interest, we removed epochs that had extreme values in any of the 9 channels of interest, in order to preserve data. An average of 22.3 trials (SD = 11.7, range = [0, 44]) were rejected from the region of interest across participants.

#### 2.3.4 Multivariate Pattern Analyses

We performed MVPA using the CoSMoMVPA toolbox ^33^ in Matlab R2017B. For each participant, a support vector machine (SVM) was trained to discriminate between the neural patterns evoked by the two semantic conditions, using the raw voltage values from all scalp electrodes and all time points. We used a standard leave-one-target-out cross-validation approach: we separated our data into a testing set containing the two trials corresponding to one target (e.g., the target “kite” in the congruent and the incongruent condition), and a training set consisting of all the other trials. We then trained an SVM algorithm to find a decision boundary that best discriminated the two experimental conditions in the training set and tested the classifier’s categorization of the two held-out trials in the testing set. We repeated this procedure 94 times, leaving a different target out each time. We then averaged the accuracy obtained for each iteration to obtain a single classifier accuracy score for each participant. If this accuracy was significantly above chance (theoretical chance level: 50%), we conclude that there was information in the brain signals that differentiated between the two semantic conditions.

To test whether the obtained accuracy was significantly above chance, we used a permutation test ^34^. This consists of randomly attributing every trial to one of the two conditions, then running the above classification procedure on these permuted data. In doing so we maintained the original target pairing (i.e., randomly swapping conditions within words) and cross-validation procedure (i.e., leaving one target word out). We repeated this procedure 1000 times to estimate a null distribution of accuracies ^34^. Accuracy was considered significant when the observed accuracy was higher than 95% of the null distributions’ accuracies (α = .05) ^34^.

We additionally wished to include an indication of effect size. In the context of multivariate decoding, this is not trivial, because standard measures of effect size fail to correct for the number of measurements and their autocorrelation as they would do in univariate analyses ^35^. This means that MVPA effect sizes calculated in a standard way may not be interpretable using the usual rules of thumb, or comparable across studies (though see ^36^ for an interpretable measure of effect size using cross-validated multivariate ANOVA). Instead, as the variance in the data is best captured by each individual’s estimated null distribution, we calculated illustrative effect sizes for each participant according to the formula:

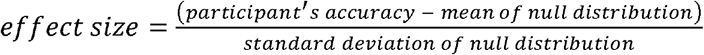

We also included an exploratory analysis to check whether MVPA detection rates would be higher if we averaged the data from several trials together before classification to create “pseudotrials”, on the basis that this might reduce noise in the data and improve classification ^37^. In order to select the best parameters for this analysis (e.g., number of trials to average to create each pseudotrial), we simulated data with a small effect using the CoSMoMVPA toolbox ^33^ and systematically tested different averaging parameters. Highest performance was found when 12 trials were averaged together in each pseudotrial. For each participant, we then analysed our experimental data by creating 100 sets of pseudotrials (each pseudotrial comprising an average of 12 real trials) and performing the classification on each of these sets as described above. Finally, we averaged the results from the 100 pseudotrials sets together to give a single measure of classification accuracy for this participant. We then repeated the same analysis using 1000 label permutations to build the null distribution for the data analysed as pseudotrials. Although this approach tended to yield higher decoding accuracy for the correctly-labelled data, it also increased the variance of the null distribution over permutations, resulting in lower sensitivity overall (8/20 participants) compared to our original approach.

Finally, we examined two possible sources of intra-individual variation on decoding accuracy: participant age and signal quality, as measured by the standardised measurement error (SME) ^38^. The SME is a newly developed measure that quantifies data quality at the individual-subject level. We obtained a measure of SME for each channel and each individual using the ERPLAB toolbox in Matlab ^39^. First, for each individual and each channel separately, we created 10,000 differential ERPs (congruent minus incongruent ERP) in a bootstrapping approach. For each bootstrap, this consisted of selecting with replacement as many congruent and incongruent epochs as the number of accepted congruent and incongruent epochs for this individual. Then, we calculated the mean amplitude compared to baseline of each differential ERP in the time-window 300 ms to 800 ms (which corresponds to the expected time-window for the N400 ^22,26,40^). Finally, we computed the SME for each channel as the standard deviation of the mean amplitude of the difference ERPs across the 10,000 bootstraps. We then investigated whether participants with noisier data (i.e., with higher SME values) had lower decoding accuracy. We first averaged the SME across all the scalp electrodes, then calculated the Spearman’s correlation coefficient between this mean SME and the individuals’ decoding accuracy.

Having established the presence of a difference in the brain signals, we then performed additional analyses to describe the timecourse and the topology of the effect. We first computed the timecourse and the topography of the MVPA results, then examined the ERP using univariate analyses of the two conditions.

#### 2.3.5 Time-resolved MVPA

In order to examine the timecourse of the discriminating brain signals, we ran a follow-up analysis where we trained the classifier to distinguish between the two conditions over time. For each time point, we trained the classifier on the data from all electrodes, at that time point and the 10 neighbouring time points (5 on each side). We then repeated this analysis across time points. To test for significance, we used a permutation test and threshold-free cluster enhancement (TFCE), as described by Smith and Nichols ^41^, on all time points excluding the baseline as in our previous work ^42^. This approach allows for extraction of a statistic of cluster level support at each time point, for the observed accuracy and the permutation results. The maximum TFCE statistic across time of each permutation was used to create a corrected null-distribution. The observed TFCE statistic at each time point was considered significant if it was larger than 95% of the null-distribution. This allowed us to observe the evolution of decoding accuracy over time, thus indicating *when* the classifier could find brain information that discriminated between conditions.

#### 2.3.6 Time-space-resolved MVPA

Finally, in order to illustrate both the timecourse and topography of the decoding, we ran a second follow-up decoding analysis allowing the classifier to use the data coming from a subset of neighbouring electrodes (5 electrodes per neighbourhood), and from a subset of neighbouring timepoints (11 timepoints per neighbourhood). We repeated this analysis for each electrode and each time point. This yielded a topographic map of decoding accuracy over time for each participant. For visualisation, we illustrate this topography for decoding accuracy averaged over time within four time-windows spanning 200 to 1000 ms post stimulus onset. Videos of the decoding accuracy evolution over time are available at https://osf.io/bv2dy/.

#### 2.3.7 Univariate analyses

We defined a region of interest centred around Cz, and the eight surrounding electrodes (Fc1, FCz, FC2, C1, C2, CP1, CPz, and CP2), based on ^26^ and ^27^.. We restricted our analyses to the data from 150 ms post-stimulus, based on previous results ^22^. We averaged the data from the nine electrodes of interest, and analysed the difference between the two conditions by running t-tests at each time point from 150 ms onward. We corrected for multiple comparisons by using a temporal cluster threshold calculated using Guthrie and Buschwald’s ^43^ method.. We first calculated the autocorrelation value of our ERP waveform. We then generated 1000 random ERP series with the same autocorrelation value as our original ERP waveform. We calculated the t-test statistics between the congruent and the incongruent condition at each timepoint for our original waveform and each of the random series. For each random series, we determined the longest run of t-values below .05. A cluster was considered significant in our original waveform if it was longer than 95% of the random series’ longest cluster. These analyses were carried out at the group level, and each individual separately.

Similarly to MVPA analyses, we used the SME as an indication of data quality, and examined whether it impacted the size of the N400 effects. To do this, we calculated the area under the difference ERP (congruent – incongruent) using a trapezoidal integration from 300 ms to 800 ms within the ROI, in steps of one sample (2 ms). These time points correspond to the expected N400 effect time course ^22,26,40^. We then calculated the Spearman correlation coefficient between this area and the mean SME for the 9 ROI channels.

### 2.4 Results

We examined children’s brain responses to semantically congruent and incongruent spoken and visual sentences using EEG. During the experiment, children simultaneously watched and listened to sentences, and were occasionally prompted to press a button to indicate whether the sentence they just heard was correct (e.g., “the squirrel stored nuts in the *tree*”) or incorrect (e.g., “the squirrel stored nuts in the *door*”).

#### 2.4.1 Behavioural results

Participants (n = 20) performed the button-press task with a high degree of accuracy (mean percent correct: M = 97.17%, SD = 3.29%, range = [90%, 100%]), indicating that they understood the meaning of the sentences and were able to notice semantic anomalies. They also responded within the required time on 99.3% of trials (mean reaction time: M = 1.63 s, SD = 0.41, range = [0.51, 4.02]).

#### 2.4.2 Multivariate Pattern Analyses results

##### 2.4.2.1 Temporally and spatially-unconstrained MVPA

Using temporally- and spatially-unconstrained multivariate classification analysis, we could decode whether the target word was semantically congruent or incongruent with the sentence in 65% of participants (13/20 participants, Figure 2). Individual decoding accuracies ranged from 68 to 49%, with effect sizes ranging from 4.08 to −.27, and p-values ranging from .0002 to .60. Thus, using all of the data available allowed us to detect statistical effects of semantic violation in two thirds of individuals. We did not find a significant correlation between decoding accuracy and participants’ age (Spearman correlation r = .136, p = .568) or the standardised measurement error across all channels (Spearman correlation r = −.36, p = .12).

**Figure 2.**
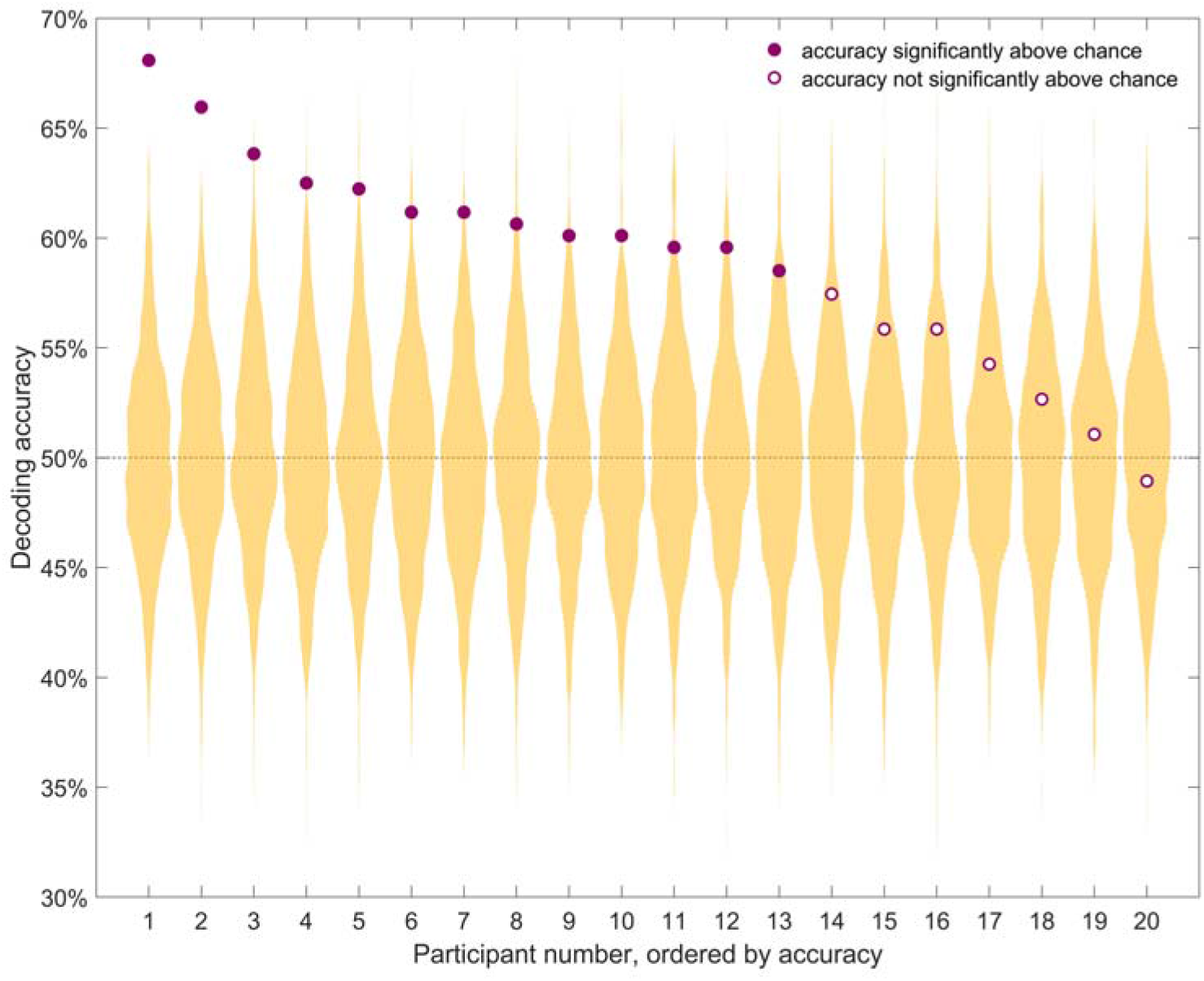
Individual decoding accuracy for classification of identical target words in congruent and incongruent contexts. Purple circles indicate decoding accuracy for each participant. The yellow distribution shows the null distribution obtained by the permutation test for that participant. Thirteen out of 20 participants had a decoding accuracy significantly above chance (full purple circles). Chance (50%) is indicated by the horizontal dashed line. Participants are sorted by decoding accuracy.

##### 2.4.2.2 Time-resolved MVPA

As our main result yields optimal statistical power by trading off spatial and temporal resolution, we conducted a series of follow-up exploratory analyses to qualify when and where the effect of interest arose. First, we used time-constrained multivariate classification analyses, to extract the temporal evolution of decoding accuracy for each participant (Figure 3). Restricting the classifier to short time windows, and repeating this over time, shows the time course with which information is decodable from the spatial pattern of activity across the scalp. This analysis revealed substantial variability in the timecourse of the effect across participants. While three participants (P5, P6 and P12) showed decodable information about the semantic condition at around 400ms, in line with classic N400 effects, others showed decodable semantic information earlier (200ms, P9) or later (P1, P3, P9). As expected, this approach was less sensitive than the main analysis, because the classifier was not given the entire length of the epoch to distinguish between the conditions, and because the multiple comparisons inherent in this approach necessitated a more stringent alpha level for inference. Of the 13 participants with significant decoding over time and space, only 7 retained enough spatial information to be decoded using this approach.

**Figure 3.**
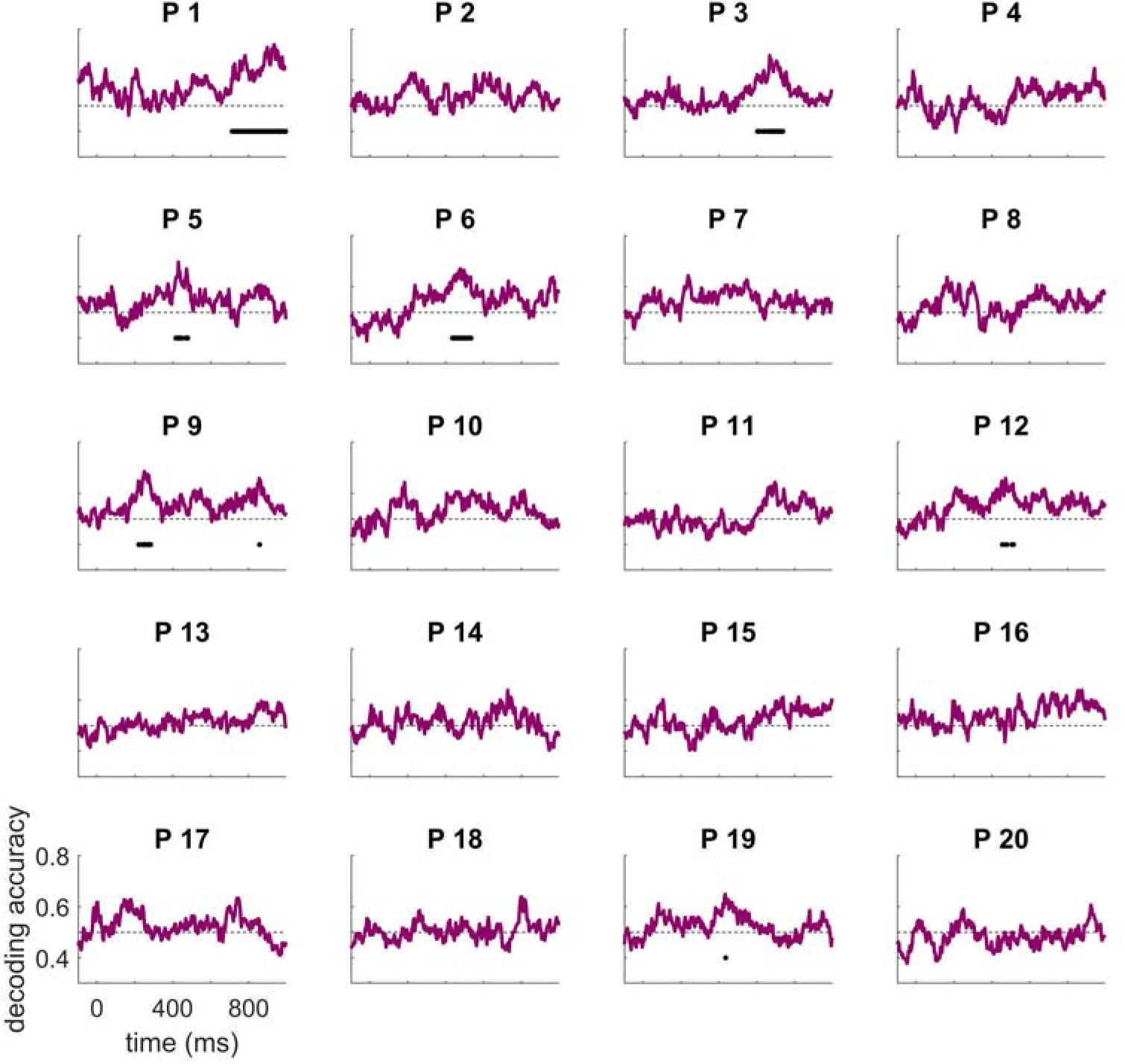
Individual time-resolved decoding accuracies. Chance (50%) is indicted by the horizontal dashed line. Accuracy significantly above chance is indicated by a solid black horizontal line for each participant (after multiple comparison correction). Participant numbering as in Figure 2, six participants had decoding accuracy significantly above chance.

##### 2.4.2.3 Time-space resolved MVPA

Next, we used a time and space resolved decoding approach to illustrate both when and where there was discriminative information for each individual (Figure 4). Although these analyses were exploratory and we did not perform significance testing, we observed substantial inter-subject variability in the regions and times that were informative for the classifier. While some participants showed high accuracy at times and regions that correspond to the typical univariate N400 effect (i.e., a centroparietal effect around 400ms, e.g., P5, P6, P8), others showed high accuracies at unexpected times (e.g., late, P11) and/or at unexpected locations (e.g., left lateralised, P9, late and occipitotemporal, P12). To further investigate these results and examine their mapping onto univariate differential responses between conditions, we additionally extracted the N400 ERP in response to the two conditions.

**Figure 4.**
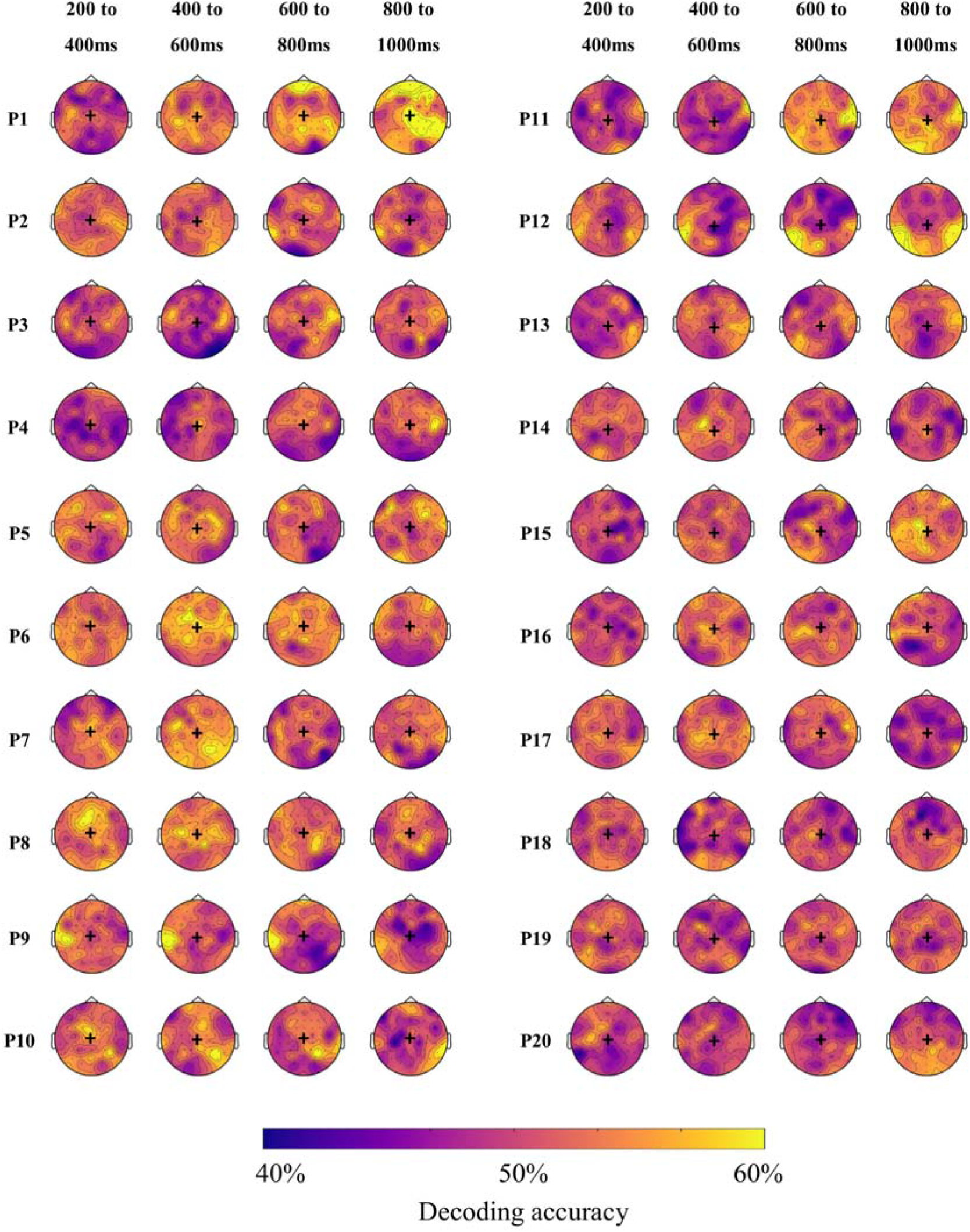
Individual topographic distribution of decoding accuracy for 200 ms time windows from 200 to 1000 ms after target onset. Yellow areas indicate higher decoding accuracy, and blue areas indicate lower decoding accuracy. These maps are for illustrative purposes and are not thresholded for significance. Black + indicates Cz location. Participant numbering as in Figure 2, participants 1-13 showed a significant effect in the main analysis.

### 2.5 Univariate analyses

Finally, we examined the N400 univariate effect by computing the voltage changes over time for the two experimental conditions (congruent and incongruent words) in a pre-specified centroparietal region of interest. This analysis was included for comparison with the wider literature and to determine the extent to which the results of our main analysis could be attributed to classic N400 effects. At the group level, we found a significant N400 effect for a cluster of timepoints from 289 – 873 ms (Figure 5, top panel). The effect was maximal at central locations from 200 – 400 ms, then extended to frontal regions at later timepoints (Figure 5, bottom panel).

**Figure 5.**
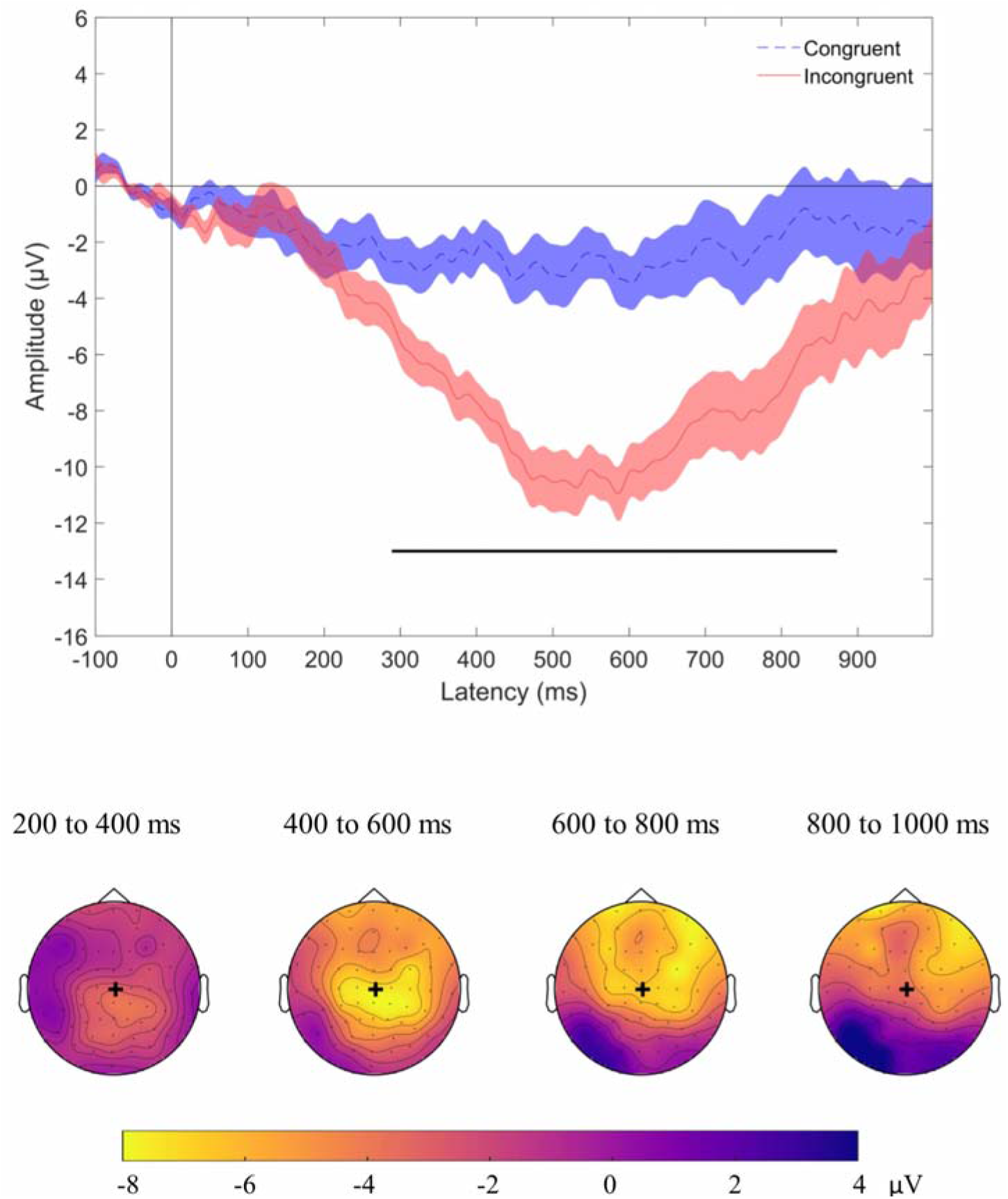
Group N400 effect. Top panel displays grand average ERPs (n=20), with congruent (dashed blue) and incongruent (solid red) conditions for the signal averaged over our region of interest. Shading indicates standard error of the mean. Time points at which there was a statistical difference between conditions are indicated with a solid black line under the plot (p<0.05, after cluster correction for multiple comparisons). Bottom panel illustrates the topographic map of the N400 effect (mean magnitude of unrelated minus related condition) from 200 to 1000 ms after target onset in the group. Yellow colours indicate a negative difference between the two conditions (incongruent more negative than congruent), while blue colours indicate a positive difference. These maps are for illustrative purposes and are not thresholded for significance. Black + indicates Cz location. The N400 effect was distributed over central and centro-frontal regions.

At the individual level, 45% of participants showed a significant univariate N400 effect for at least one significant cluster of time points in our region of interest (9/20 participant, figure 6). To summarise across analyses, 7 of the 13 participants with significant decoding results also showed a detectable N400 effect, while the remaining 5 did not. Only one participant (P17) showed a detectable N400 effect in the absence of significant MVPA classification.

**Figure 6.**
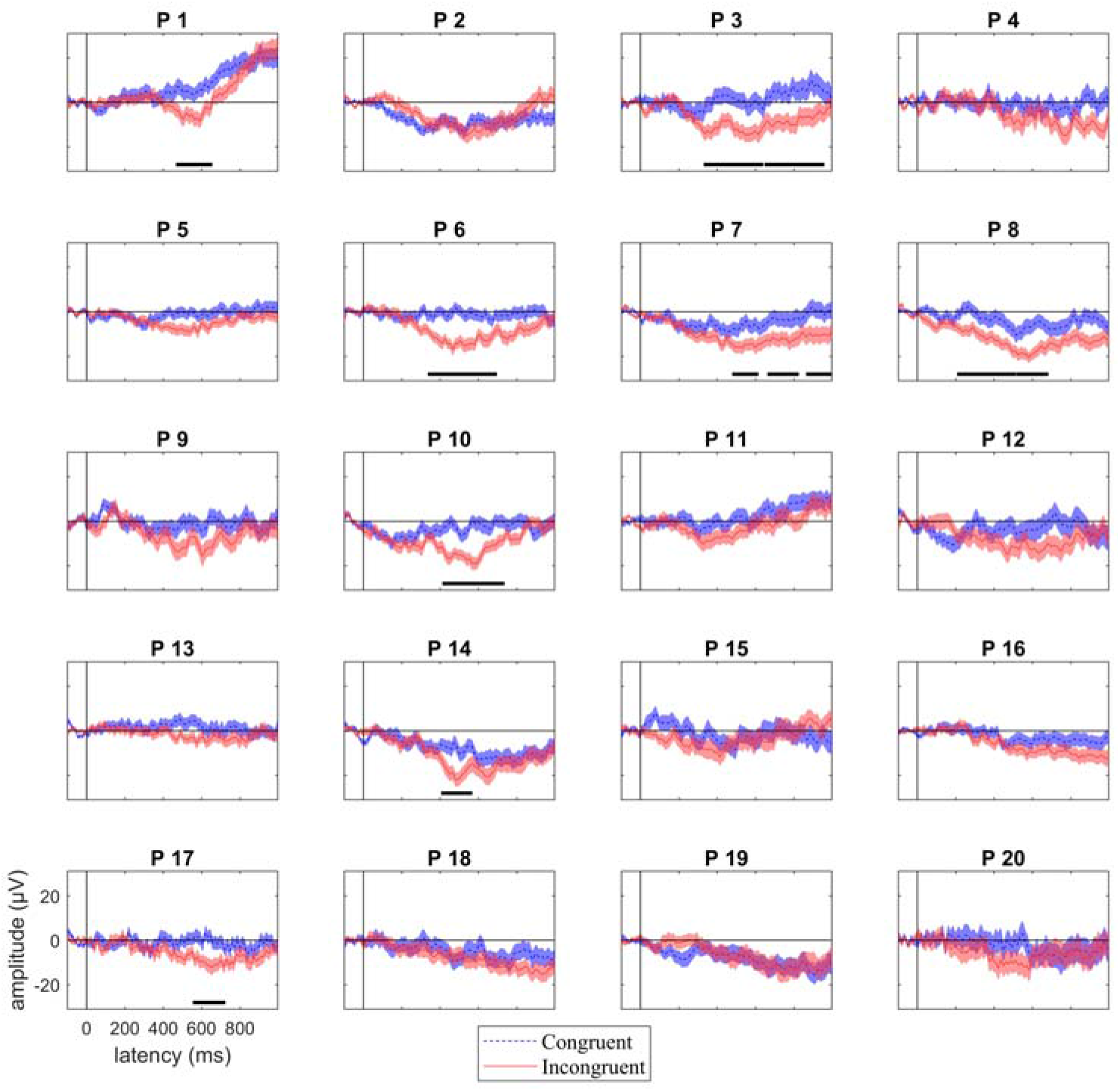
Individual participant responses to target words following congruent (dashed blue) and incongruent (solid red) sentences in our region of interest. Shading indicates standard error of the mean. Time points where there was a statistically significant N400 effect are indicated with a solid, horizontal, black line. P indicates participant, with participant numbering as in Figure 2. 45% (9/20) of participants showed a statistical univariate N400 effect.

In addition, we did not find a significant correlation between the amplitude of the N400 effect and data quality, as indicated by the standardised measurement error (Spearman’s rho = −.30, p =.19).

The topology of the N400 effect was highly variable across participants (Figure 7) in line with the time-space-resolved MVPA analysis (above) and our previous work ^27^.

**Figure 7.**
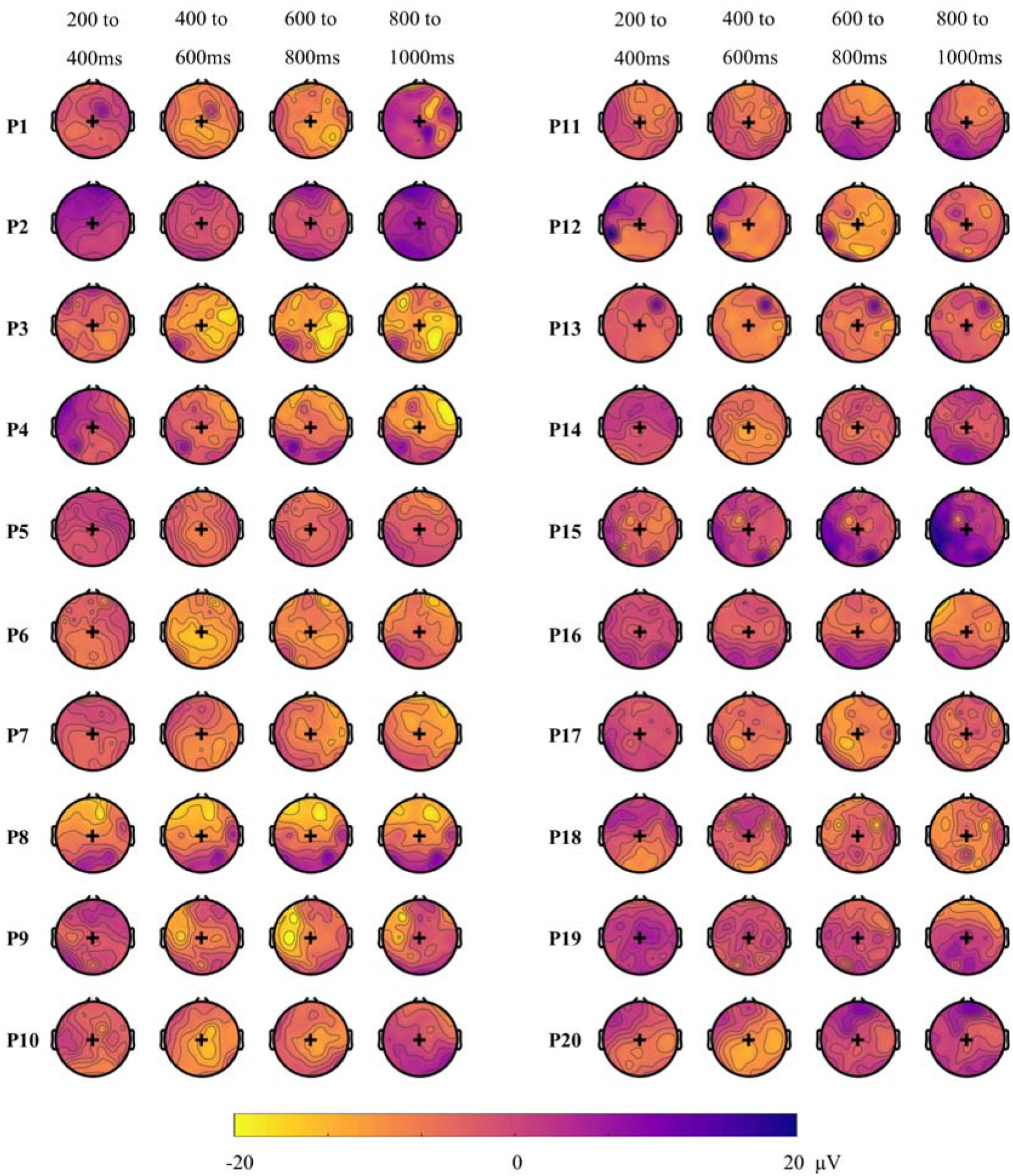
Individual topographic maps of the N400 effect (unrelated minus related condition) for 200 ms time windows from 200 to 1000 ms after target onset. Yellow colours indicate a negative difference between the two conditions (incongruent more negative than congruent), while blue colours indicate a positive difference. These maps are for illustrative purposes and are not thresholded for significance. Back + indicates Cz location. The topography of the N400 effect varied across individuals. P indicates participant, with participant numbering as in Figure 2.

## 3 Discussion

Recent electrophysiological evidence has shed light on the semantic processing of spoken language in minimally-verbal autistic children ^25^. This has important implications for our understanding of brain processing in autism, and for the care and treatment of these individuals. However, heterogeneity in the neural signals, even in neurotypical children, is yet to be addressed. This study validates a multi-modal N400 paradigm to assess lexico-semantic processing from electrophysiological activity, and reports on the reliability of neural signals across individual neurotypical children. We recorded EEG from children while they were watching video-animated sentences with matched correct and incorrect endings, and used two complementary approaches to analyse their brain data. Using Multivariate Pattern Analyses (MVPA) to pool information over both space and time, we detected patterns of brain activity that discriminated between the two semantic conditions in 65% of individual children. Further descriptive analyses suggested that the patterns of discriminative activity were variable across individuals, ranging both in topography and in time. We additionally analysed the N400 ERP using a univariate approach, and found a robust N400 effect in the central location at the group level, as well as in 45% of individual participants. We present a summary of these analyses in Table 2.

**Table 2.** Summary of individuals’ effect in different analyses. ‘Yes’ indicates that we found a significant effect for this participant.

| Participant number | Time- and space-<br>unconstrained MVPA | Time-resolved MVPA | Univariate N400 |
| --- | --- | --- | --- |
| 1 | *Yes* | *Yes* | *Yes* |
| 2 | *Yes* | No | No |
| 3 | *Yes* | *Yes* | *Yes* |
| 4 | *Yes* | No | No |
| 5 | *Yes* | *Yes* | *Yes* |
| 6 | *Yes* | *Yes* | *Yes* |
| 7 | *Yes* | No | *Yes* |
| 8 | *Yes* | No | *Yes* |
| 9 | *Yes* | *Yes* | No |
| 10 | *Yes* | No | *Yes* |
| 11 | *Yes* | No | No |
| 12 | *Yes* | *Yes* | No |
| 13 | *Yes* | No | *Yes* |
| 14 | No | No | No |
| 15 | No | No | No |
| 16 | No | No | No |
| 17 | No | No | *Yes* |
| 18 | No | No | No |
| 19 | No | *Yes* | No |
| 20 | No | No | No |
| **Total** | 13/20 | 7/20 | 9/20 |

Our data replicate recent findings that univariate effects to identical auditory tokens presented in different semantic contexts can be reliably observed in about half of the participants ^26,42^. This result has important implications both for clinical application ^23,24,40^ and for researchers using the N400 to study language acquisition ^44,45^.For the participants not showing reliable differences, it is unclear whether our methods were not sensitive enough to detect N400 effects to semantic violations, or whether N400 effects were truly absent in some participants. In the latter case, it remains unclear whether our failure to detect differential brain responses in some participants was due to their reliance on different cognitive processes, or whether similar cognitive processes were supported by different neural substrates. It is also possible that for some participants, the incongruent sentences were not eliciting strong semantic or predictive violations, or that these violations became less strong over the course of the experiment. In an attempt to pinpoint the possible cause of inter-individual variability, we examined the impact of data quality on the neural responses recorded at the individual-subject level. Although data quality will obviously affect our ability to detect meaningful effects, variation in the standardised measurement error (SME) did not account for variation in decoding accuracy or the amplitude of the univariate difference wave across subjects. The SME was recently introduced as a universal measure of data quality in individuals, and reflects the standard deviation of a given measure, in our case the amplitude of the difference ERP waveform (congruent minus incongruent) across trials. Although we did not find evidence that lower SME was associated with larger effects in our sample, we must be cautious in interpreting correlation coefficients drawn from a relatively small sample (20 participants). Future studies may benefit from larger sample sizes, or from including alternative measures of signal-to-noise ratio, for example by examining the amplitude of auditory evoked responses to simple tones ^46,47^ or examining neural entrainment by speech envelopes ^48,49^. We also did not find any evidence for an impact of age on our effects, but again with only 20 participants we must be cautious about interpreting this absence of correlation. It would be interesting to assess other potential sources of inter-individual differences, such as language ability or lateralisation, on a larger sample.

Previous work using purely auditory paradigms have found a similar ^26,27,exp 1^ or better ^26,27,exp 2^ detection rate of brain responses to semantic violations than the current study. There was therefore no evidence that adding a visual component made the N400 effect stronger in individuals. We hypothesized that creating a semantic context using both visual and auditory information would increase the participants’ expectation for the final word, making an incongruent ending even less predictable. This prediction was partly based on previous findings that the N400 reflects lexico-semantic access irrespective of the modality. For example, Nigam et al. found similar N400 effects for final words of written sentences, whether the final words were in written or picture format ^50^. Similarly, when testing the N400 in response to auditory words versus written words, Holcomb and Neville found overlapping neural processes between these two modalities ^51^. However, in our study, it is also possible that adding a visual component had a detrimental effect, for example by shifting attention away from listening to the sentences. The high accuracy of participants on the task suggests that participants were able to correctly classify the congruency of the sentences when probed, but this is not a sensitive measure of attention. In the future, presenting the animations before the auditory sentences, instead of concurrently, may alleviate this concern. It is also possible that our numerically lower sensitivity to detect discriminative neural patterns compared to our previous study, with a similar stimuli and identical analyses, was due to the task that participants performed. In Petit et al. ^27^, participants had to count the number of incorrect sentences within a block. Here, participants were occasionally probed for whether the last sentence they heard was correct. Although the probing task is attractive for its simplicity (minimal working memory component and no requirement to be able to count), it is possible that the counting task in ^27^ encouraged more sustained attention to all the stimuli, potentially improving decoding accuracy. However, the two studies are not directly comparable because the current study had more unique sentences and less repetition(188 trials unique sentences presented once in this study versus 112 unique sentences presented twice for a total of 224 trials in ^27^), and because the current study and examined a smaller age range (9 – 12 years, versus. 6-12 year in ^27^). There were also differences in the way the sentences for the two studies were recorded an presented: here complete sentences were recorded and spliced, with adjectives added to a subset of the original sentences to introduce a plosive for splicing, whereas in ^27^ sentence frames were recorded separately from target words resulting in penultimate word lengthening and were presented with a 100 ms silent gap before the target.

Using MVPA allowed us to detect differential brain responses to congruent and incongruent sentences in 65% of individuals (13/20 participants). Of these 13 only 8 (62%) also showed a significant N400 effect. Of the 7 participants who did not have a significant classification result, only one showed a significant univariate N400 effect. This highlights the higher sensitivity of MVPA to detect differences in the pattern of brain activity that are relevant to the semantic condition but may occur at variable times or locations, or as subtle changes in the pattern of brain activation, across individuals. In our implementation, MVPA takes data from all sensors and timepoints into account without introducing multiple comparisons (only one statistical test is performed per participant). In our univariate approach we minimised multiple comparisons by restricting our analyses to a pre-defined ROI, but with the trade-off of not being sensitive to potential differences occurring outside of this region. Only two studies have investigated decoding accuracy to classify individuals’ EEG data. Geuze et al. have used a similar approach as the current study on adult participants who were presented with words that were related or unrelated with a previously-heard probe word and found above-chance decoding for all of their participants ^52^. It is likely that their higher accuracy reflects the better data quality recorded in adults, who are better able to sit still and engage with the task. More recently, Tanaka et al. reported significant decoding of single-trial semantic anomalies, with an overall accuracy of 59.5% ^53^. They, however, did not report individual-participant accuracies, making it hard to compare their results to the current findings. The current study adds to this body of literature by illustrating the capacity of classifiers to detect processing of semantic congruency in individual children performing a covert task, using rigorous and robust EEG data analyses.

Although MVPA appears to be more sensitive for detecting differences in brain activation between conditions, it does not always perform better than univariate analyses. For one participant in our study (P 17), decoding accuracy was not significantly different from chance, while the N400 univariate effect was statistically significant. At least two accounts may explain this surprising result. Firstly, MVPA was performed on all the trials, while we cleaned the data during preprocessing before univariate analyses. It is possible that the classifier could not overlook the noise in the data or that the univariate effect was driven by a few trials, and was not consistent enough to be detected as a pattern by the classifier. We chose not to clean the data before performing MVPA as our leave-one-target out approach requires the dataset to be balanced, with the same trials present in both conditions. Removing noisy trials would have involved removing their counterpart in the other condition, as well as the trials in both conditions that had the same sentence frame as the noisy trials. Thus, for each noisy trial, we would have to reject four trials in order to keep a balanced design, which would leave too few trials to use for classification. Secondly, because the N400 effect occurred at the expected location and timing in that individual, it is likely that the univariate approach was more sensitive. However, for the purpose of a neural test of language comprehension in non-communicative populations, this typicality cannot be assumed, so it may nonetheless be preferable to use methods that can detect differences in brain activity at unexpected locations and timing. This is reflected in our data, where five participants had significant decoding but did not show an N400 effect at our pre-specified region of interest. Participant 4, for example, had a strong decoding result but showed an N400-like effect only at antero-frontal locations (see figure 7), which would not be detected with conventional N400 analyses.

## 4 Conclusion

This study aimed to assess the reliability of neural signals in response to semantic anomalies in individual neurotypical children. We used a paradigm contrasting congruent and incongruent sentences, and analysed the EEG response using different methods to illustrate the heterogeneity of children’s brain responses. Despite the challenges associated with analysing individuals’ EEG data, we were able to reliably detect neural responses to lexico-semantic anomalies in 65% of individual children using MVPA. Out of these, only a subset (62% of these) also showed a typical N400 ERP at the central location, indicating substantial individual variability in the neural basis of lexical-semantic processing. For the purpose of assessing language comprehension in special populations, such as non-verbal children with autism, this paradigm yields medium sensitivity. In a clinical setting, we would recommend to use the information from multiple types of analyses. The different analyses that we present here explore the same process of lexico-semantic decisions with different assumptions about the information recorded from the brain. By comparing multiple sources of information, we can achieve higher accuracy in the detection of semantic processing: in participants showing both decoding and a significant N400 effect, we have strong evidence for intact semantic processing, constituting an important step in designing a neural test of language comprehension.

## 6 Author contributions

Conceptualisation: S.P., A.W., N.A.B; Stimulus and task development: S.P., A.W; Data acquisition: S.P.; Formal analysis: S.P., A.W., T.G.; Writing (original draft): S.P; Writing (reviewing and editing): all authors; Supervision: A.W.

## 7 Acknowledgements

This work was funded by a Macquarie University Cognitive Science Postgraduate Research Grant to S.P and a Cross Programme Support Scheme grant from the ARC Centre of Excellence in Cognition and its Disorders to A.W. and N.B. S.P was supported by an international Research Training Program scholarship from Macquarie University. A.W. was supported by an ARC Future Fellowship (FT170100105) and MRC intramural funding SUAG/052/G101400. We thank Gabriella Keys for drawing and producing the animations.

## 9 Data availability

All raw EEG data is available at https://osf.io/bv2dy/.

## References

1. Chan, A. S., Cheung, J., Leung, W. W. M., Cheung, R. & Cheung, M. Verbal Expression and Comprehension Deficits in Young Children With Autism. Focus Autism Dev. Disabil. 20, 117–124 (2005).

2. Kjelgaard, M. M. & Tager-Flusberg, H. An investigation of language impairment in autism: Implications for genetic subgroups. Lang. Cogn. Process. 16, 287–308 (2001).

3. Sigurdardottir, S. & Vik, T. Speech, expressive language, and verbal cognition of preschool children with cerebral palsy in Iceland. Dev. Med. Child Neurol. 53, 74–80 (2011).

4. Giacino, J. T. & Smart, C. M. Recent advances in behavioral assessment of individuals with disorders of consciousness: Curr. Opin. Neurol. 20, 614–619 (2007).

5. Geytenbeek, J. et al. Utility of language comprehension tests for unintelligible or non-speaking children with cerebral palsy: a systematic review. Dev. Med. Child Neurol. 52, e267–277 (2010).

6. Tager-Flusberg, H. & Kasari, C. Minimally Verbal School-Aged Children with Autism Spectrum Disorder: The Neglected End of the Spectrum. Autism Res. 6, 468–478 (2013).

7. Kasari, C., Brady, N., Lord, C. & Tager-Flusberg, H. Assessing the minimally verbal school-aged child with autism spectrum disorder. Autism Res. Off. J. Int. Soc. Autism Res. 6, 479–493 (2013).

8. Plesa Skwerer, D., Jordan, S. E., Brukilacchio, B. H. & Tager-Flusberg, H. Comparing methods for assessing receptive language skills in minimally verbal children and adolescents with autism spectrum disorders. Autism 20, 591–604 (2016).

9. Cruse, D. et al. Bedside detection of awareness in the vegetative state: a cohort study. The Lancet 378, 2088–2094 (2011).

10. Owen, A. M. & Coleman, M. R. Detecting awareness in the vegetative state. Ann. N. Y. Acad. Sci. 1129, 130–138 (2008).

11. Harrison, A. H. & Connolly, J. F. Finding a way in: a review and practical evaluation of fMRI and EEG for detection and assessment in disorders of consciousness. Neurosci. Biobehav. Rev. 37, 1403–1419 (2013).

12. Hinterberger, T., Birbaumer, N. & Flor, H. Assessment of cognitive function and communication ability in a completely locked-in patient. Neurology 64, 1307 (2005).

13. Kotchoubey, B. et al. Information processing in severe disorders of consciousness: Vegetative state and minimally conscious state. Clin. Neurophysiol. 116, 2441–2453 (2005).

14. Mathalon, D. H., Faustman, W. O. & Ford, J. M. N400 and automatic semantic processing abnormalities in patients with schizophrenia. Arch. Gen. Psychiatry 59, 641–648 (2002).

15. Sharma, A. et al. Abnormal N400 Semantic Priming Effect May Reflect Psychopathological Processes in Schizophrenia: A Twin Study. Schizophr. Res. Treat. 2017, (2017).

16. Coderre, E. L., Chernenok, M., Gordon, B. & Ledoux, K. Linguistic and Non-Linguistic Semantic Processing in Individuals with Autism Spectrum Disorders: An ERP Study. J. Autism Dev. Disord. 47, 795–812 (2017).

17. Pijnacker, J., Geurts, B., van Lambalgen, M., Buitelaar, J. & Hagoort, P. Exceptions and anomalies: An ERP study on context sensitivity in autism. Neuropsychologia 48, 2940–2951 (2010).

18. Wang, S., Yang, C., Liu, Y., Shao, Z. & Jackson, T. Early and late stage processing abnormalities in autism spectrum disorders: An ERP study. PloS One 12, e0178542 (2017).

19. Sinclair, J. Why I dislike “person first” language. Auton. Crit. J. Interdiscip. Autism Stud. 1, (2013).

20. Kenny, L. et al. Which terms should be used to describe autism? Perspectives from the UK autism community. Autism Int. J. Res. Pract. 20, 442–462 (2016).

21. Gernsbacher, M. A. Editorial Perspective: The use of person-first language in scholarly writing may accentuate stigma. J. Child Psychol. Psychiatry 58, 859–861 (2017).

22. Kutas, M. & Federmeier, K. D. Thirty years and counting: finding meaning in the N400 component of the event-related brain potential (ERP). Annu. Rev. Psychol. 62, 621–647 (2011).

23. Cantiani, C. et al. From Sensory Perception to Lexical-Semantic Processing: An ERP Study in Non-Verbal Children with Autism. PLOS ONE 11, e0161637 (2016).

24. Coderre, E. L. et al. Implicit Measures of Receptive Vocabulary Knowledge in Individuals With Level 3 Autism. Cogn. Behav. Neurol. 32, 95 (2019).

25. DiStefano, C., Senturk, D. & Spurling Jeste, S. ERP Evidence of Semantic Processing in Children with ASD. Dev. Cogn. Neurosci. 100640 (2019) doi:10.1016/j.dcn.2019.100640.

26. Cruse, D. et al. The reliability of the N400 in single subjects: Implications for patients with disorders of consciousness. NeuroImage Clin. 4, 788–799 (2014).

27. Petit, S. et al. Towards an individualised neural assessment of receptive language in children. bioRxiv 566752 (2020) doi:10.1101/566752.

28. Block, C. K. & Baldwin, C. L. Cloze probability and completion norms for 498 sentences: Behavioral and neural validation using event-related potentials. Behav. Res. Methods 42, 665–670 (2010).

29. van Heuven, W. J. B., Mandera, P., Keuleers, E. & Brysbaert, M. SUBTLEX-UK: a new and improved word frequency database for British English. Q. J. Exp. Psychol. 2006 67, 1176–1190 (2014).

30. Delorme, A. & Makeig, S. EEGLAB: an open source toolbox for analysis of single-trial EEG dynamics including independent component analysis. J. Neurosci. Methods 134, 9–21 (2004).

31. Acunzo, D. J., Mackenzie, G. & van Rossum, M. C. W. Systematic biases in early ERP and ERF components as a result of high-pass filtering. J. Neurosci. Methods 209, 212–218 (2012).

32. Tanner, D., Morgan-Short, K. & Luck, S. J. How inappropriate high-pass filters can produce artifactual effects and incorrect conclusions in ERP studies of language and cognition. Psychophysiology 52, 997–1009 (2015).

33. Oosterhof, N. N., Connolly, A. C. & Haxby, J. V. CoSMoMVPA: Multi-Modal Multivariate Pattern Analysis of Neuroimaging Data in Matlab/GNU Octave. Front. Neuroinformatics 10, 27 (2016).

34. Maris, E. & Oostenveld, R. Nonparametric statistical testing of EEG- and MEG-data. J. Neurosci. Methods 164, 177–190 (2007).

35. Hebart, M. N. & Baker, C. I. Deconstructing multivariate decoding for the study of brain function. NeuroImage 180, 4–18 (2018).

36. Allefeld, C. & Haynes, J.-D.Searchlight-based multi-voxel pattern analysis of fMRI by cross-validated MANOVA. NeuroImage 89, 345–357 (2014).

37. Grootswagers, T., Wardle, S. G. & Carlson, T. A. Decoding dynamic brain patterns from evoked responses: A tutorial on multivariate pattern analysis applied to time-series neuroimaging data. J. Cogn. Neurosci. 29, 677–697 (2017).

38. Luck, S. J., Stewart, A. X., Simmons, A. M. & Rhemtulla, M. Standardized Measurement Error: A Universal Measure of Data Quality for Averaged Event-Related Potentials (v20b). https://osf.io/dwm64 (2020) doi:10.31234/osf.io/dwm64.

39. Lopez-Calderon, J. & Luck, S. J. ERPLAB: an open-source toolbox for the analysis of event-related potentials. Front. Hum. Neurosci. 8, (2014).

40. Beukema, S. et al. A hierarchy of event-related potential markers of auditory processing in disorders of consciousness. NeuroImage Clin. 12, 359–371 (2016).

41. Smith, S. M. & Nichols, T. E. Threshold-free cluster enhancement: addressing problems of smoothing, threshold dependence and localisation in cluster inference. NeuroImage 44, 83–98 (2009).

42. Petit, S. et al. Towards an individualised neural assessment of receptive language in children. bioRxiv 566752 (2019) doi:10.1101/566752.

43. Guthrie, D. & Buchwald, J. S. Significance testing of difference potentials. Psychophysiology 28, 240–244 (1991).

44. Borovsky, A., Elman, J. L. & Kutas, M. Once is Enough: N400 Indexes Semantic Integration of Novel Word Meanings from a Single Exposure in Context. Lang. Learn. Dev. Off. J. Soc. Lang. Dev. 8, 278–302 (2012).

45. Rämä, P., Sirri, L. & Serres, J. Development of lexical–semantic language system: N400 priming effect for spoken words in 18- and 24-month old children. Brain Lang. 125, 1–10 (2013).

46. Mair, I. W. S. & Laukli, E. Identification of Early Auditory-Evoked Responses. Audiology 19, 384–394 (1980).

47. Bieber, R. E., Fernandez, K., Zalewski, C., Cheng, H. & Brewer, C. C. Stability of Early Auditory Evoked Potential Components Over Extended Test-Retest Intervals in Young Adults. Ear Hear. Publish Ahead of Print, (9000).

48. Horton, C., Srinivasan, R. & D’Zmura, M. Envelope responses in single-trial EEG indicate attended speaker in a `cocktail party’. J. Neural Eng. 11, 046015 (2014).

49. O’Sullivan, J. A. et al. Attentional Selection in a Cocktail Party Environment Can Be Decoded from Single-Trial EEG. Cereb. Cortex 25, 1697–1706 (2015).

50. Nigam, A., Hoffman, J. E. & Simons, R. F. N400 to Semantically Anomalous Pictures and Words. J Cogn. Neurosci. 4, 15–22 (1992).

51. Holcomb, P. J. & Neville, H. J. Auditory and Visual Semantic Priming in Lexical Decision: A Comparison Using Event-related Brain Potentials. Lang. Cogn. Process. 5, 281–312 (1990).

52. Geuze, J., Gerven, M. A. J. van, Farquhar, J. & Desain, P. Detecting Semantic Priming at the Single-Trial Level. PLOS ONE 8, e60377 (2013).

53. Tanaka, H., Watanabe, H., Maki, H., Sakriani, S. & Nakamura, S. Electroencephalogram-Based Single-Trial Detection of Language Expectation Violations in Listening to Speech. Front. Comput. Neurosci. 13, (2019).

