## Supplementary table 1 for "Unconstrained multivariate EEG decoding can help detect lexical-semantic processing in individual children"

### Supplementary Table 1: List of stimuli

| **Sentence frame** | **Congruent target** | **Incongruent target** |
| --- | --- | --- |
| While eating Steve accidentally bit his | tongue | kite |
| It was windy enough to fly a | kite | tongue |
| The directions did not match any roads on the old | map | pace |
| Jessie ran the race at a slower | pace | map |
| At the royal wedding the princess married the | prince | chair |
| The man sat down on the comfortable | chair | prince |
| For her school dance Becky wore a new | dress | bridge |
| The boat passed easily under the | bridge | dress |
| The little girl was very afraid of the | dark | bread |
| She went to the bakery to buy a loaf of | bread | dark |
| The dentist says to brush your teeth twice a | day | band |
| She played the guitar so she joined the | band | day |
| The student went to the library to read a | book | paint |
| He wanted color in the room so he bought a tin of | paint | book |
| The squirrel stored nuts in the | tree | bike |
| A good way to exercise is to ride a | bike | tree |
| During the lecture Jen kept checking the | time | queen |
| The princess may someday become a | queen | time |
| After dinner they washed the | dishes | trees |
| In autumn leaves fall off the | trees | dishes |
| Most students prefer to work during the | day | tea |
| I put the kettle on to make a hot cup of | tea | day |
| After pulling him over, the police officer told the man to get out of his | car | book |
| He turned the page of his favorite | book | car |
| She forgot her watch so she asked for the | time | book |
| He read a chapter of the | book | time |
| She got out of the car and closed the | door | cake |
| There were candles on the birthday | cake | door |
| I prefer cats rather than | dogs | teeth |
| The dentist opened her mouth to check her | teeth | dogs |
| For his wife's birthday he baked a | cake | time |
| Her job was easy most of the | time | cake |
| At night the old woman locked the | door | break |
| Several hours into his shift Lyle was ready for a | break | door |
| During the winter holidays people tend to eat a lot of good | food | door |
| I had no key to open the | door | food |
| His father was the coach of the football | team | door |
| After coming inside Bob locked the | door | team |
| The indoor plant was growing bigger and needed a new | pot | broom |
| John swept the floor with a | broom | pot |
| The moon was shining in the night | sky | vase |
| Katie put the flowers in a big | vase | sky |
| She put on her sunglasses to protect her tired | eyes | marks |
| Due to his hard work he received excellent | marks | eyes |
| The surfer is scared of being bitten by a great white | shark | ink |
| The cheap pen quickly ran out of black | ink | shark |
| The dragon was slain by the valiant | knight | stick |
| To help her walk around the girl used a walking | stick | knight |
| Father carved the turkey with a sharp | knife | ink |
| The pen in his pocket had unfortunately leaked | ink | knife |
| He washed his hands with water and | soap | knife |
| At dinner he cut his steak with a sharp | knife | soap |
| He had a long day and was in a bad | mood | splash |
| She jumped in the pool and made a big | splash | mood |
| To keep the dogs out of the yard he built a six foot | fence | snack |
| Before practice he decided to eat a quick | snack | fence |
| She made him a sandwich for his packed | lunch | cell |
| The prisoner fell asleep inside his dark | cell | lunch |
| She looked up at night to see a sky full of bright | stars | aim |
| Tom’s arrows missed the target due to his bad | aim | stars |
| For breakfast he ate scrambled | eggs | socks |
| Derek’s feet were cold so he put on some thick | socks | eggs |
| Cid needed a belt to hold up his | pants | teeth |
| After every meal it’s good to brush your | teeth | pants |
| His shirt was so worn it had a big | hole | walk |
| The old man has to use a cane to go on a short | walk | hole |
| The genie granted the man his first | wish | sky |
| The stars are high up in the night | sky | wish |
| There were no extra seats so she sat on the concrete | floor | milk |
| The little girl left Santa a plate of cookies and | milk | floor |
| She could tell he was mad by the tone of his loud | voice | milk |
| The hungry baby wanted to drink | milk | voice |
| He went to the lake to catch a big | fish | wall |
| She hung the painting up on the white | wall | fish |
| He took his dog out for a short | walk | sun |
| The baseball player’s cap protected him from the hot | sun | walk |
| I roasted the marshmallow over the hot | fire | ring |
| Bob proposed and gave her a diamond | ring | fire |
| It was dark in the room so she turned on the bright | light | ring |
| The man presented his new fiancée with an expensive diamond | ring | light |
| Sarah saw animals from around the world at the country's best | zoo | sword |
| The knight got ready for battle and drew his magnificent | sword | zoo |
| To cut the chicken Sue needed a sharp | knife | farm |
| They raised pigs on their private | farm | knife |
| Tim joined the airforce because he always wanted to fly a | plane | desk |
| To do her homework she sat down at her | desk | plane |
| The real estate agent sold the big | house | word |
| Today the baby spoke his first | word | house |
| Most cats see very well at | night | floor |
| I accidentally spilled some water on the concrete | floor | night |
| After driving for hours Kevin rested at the next | stop | hand |
| When he touched the hot stove the child burned his right | hand | stop |
| Pete broke his arm and needed to wear a | cast | pan |
| Jane fried some bacon in the | pan | cast |
